# DNA-mediated assembly of multi-specific antibodies for T cell engaging and tumor killing

**DOI:** 10.1101/782862

**Authors:** Liqiang Pan, Chan Cao, Changqing Run, Liujuan Zhou, James J. Chou

## Abstract

Targeting T-cells against cancer cells is a direct means of treating cancer, and already showed great responses in clinical treatment of B-cell malignancies. A simple way to redirect T-cells to cancer cells is using multi-specific antibody (MsAb) that contains different arms for specifically “grabbing” the T-cells and cancer cells; as such, the T-cells are activated upon target engagement and the killing begins. Here, a Nucleic Acid mediated Protein-Protein Assembly (NAPPA) approach is implemented to construct a MsAb for T-cell engaging and tumor killing. Anti -CD19 and -CD3 single-chain variable fragments (scFvs) each are conjugated to different L-DNAs with sequences that form the Holliday junction, thus allowing spontaneous assembly of homogeneous protein-DNA oligomers containing two anti-CD19 and one anti-CD3 scFvs. The new MsAb shows strong efficacy in inducing Raji tumor cell cytotoxicity in the presence of T-cells with EC_50_ ~ 0.2 nM; it also suppresses tumor growth in the Raji xenograft mouse model. The data indicate that MsAbs assembled from protein-DNA conjugates are effective macromolecules for directing T-cells for tumor killing. The modular nature of the NAPPA platform allows rapid generation of complex MsAbs from simple antibody fragments, while offering a general solution for preparing antibodies with high-order specificity.

---

The essence of bispecific antibody (BsAb) is that two different antigen binding domains are physically linked such that they can engage multiple cells presenting different antigens, and many creative methods have been developed to achieve this property^1, 2^. In many of the early proof-of-concept studies, BsAbs were generated by chemically crosslinking two different IgGs or Fabs using bifunctional crosslinking reagents that react specifically with the thiol and primary amine groups of the antibody^3, 4^. Although several of BsAbs prepared in this way have advanced to clinical trials^5–7^, large majority of BsAbs currently being developed are generated with recombinant antibody engineering. Over 100 different formats of multi-specific antibodies (MsAbs) have been engineered based on the immunoglobulin G (IgG) or its components (reviewed in Ref ^1^), some containing the Fc and others do not. Well-known examples of Fc-less formats are tandem single-chain variable fragments (scFvs)^8^ and tandem nanobodies^9^. Among them, the bispecific T-cell engager (BiTE) consisting of tandem anti-CD19 and anti-CD3 scFvs (blinatumomab) is the first FDA approved BsAb, used for treating acute lymphoblastic leukemia^10, 11^. Natural IgG containing the Fc is symmetric. To introduce bispecific or asymmetric property to the IgG, a variety of methods have been developed to favor heterodimeric heavy chain pairing. A few prominent examples are knob-into-hole^12^, structural-based mutagenesis^13^, and electrostatic steering^14^ that favors heterodimerization or disfavors homodimerization of the Fc. Further, combining the knob-into-hole and appending one of the two arms of the IgG with another scFv or Fab allows the assembly of trispecific antibody^15^.

The above are only a few examples of BsAb engineering. The fact that many of the BsAbs with different sizes and forms all showed strong ability to mediate T-cell engaging suggests that if the antibody domains targeting different antigens stay intact within a molecular framework, the form of the framework may not be so important as long as it does not place the antigen-recognition domains too far apart.

Inspired by the previous works, we implemented a <u>n</u>ucleic <u>a</u>cid mediated <u>p</u>rotein-<u>p</u>rotein <u>a</u>ssembly method^16^, designated NAPPA, to construct MsAbs to drive T-cell – tumor cell engagement and tested the *in vitro* and *in vivo* efficacy of the new molecular complexes. In this implementation (illustrated in **Figure 1**), scFvs targeting different antigens each were chemically conjugated, at their C-termini, to a 30-base DNA that was designed to pair with DNAs linked to other scFvs, and the scFv-DNA conjugates were purified separately. Hence, in this format, the scFvs are the antigen binding modules (ABMs) and DNAs are the assembly modules. We could then assemble the multi-specific scFv oligomers by simply mixing the pre-purified and -stored scFv-DNAs at the designated molar ratio. Owing to the accuracy of DNA base pairing interactions, the spontaneously assembled scFv-DNA oligomers should be homogeneous, stable, and readily usable in T-cell engaging experiments.

**Figure 1.**
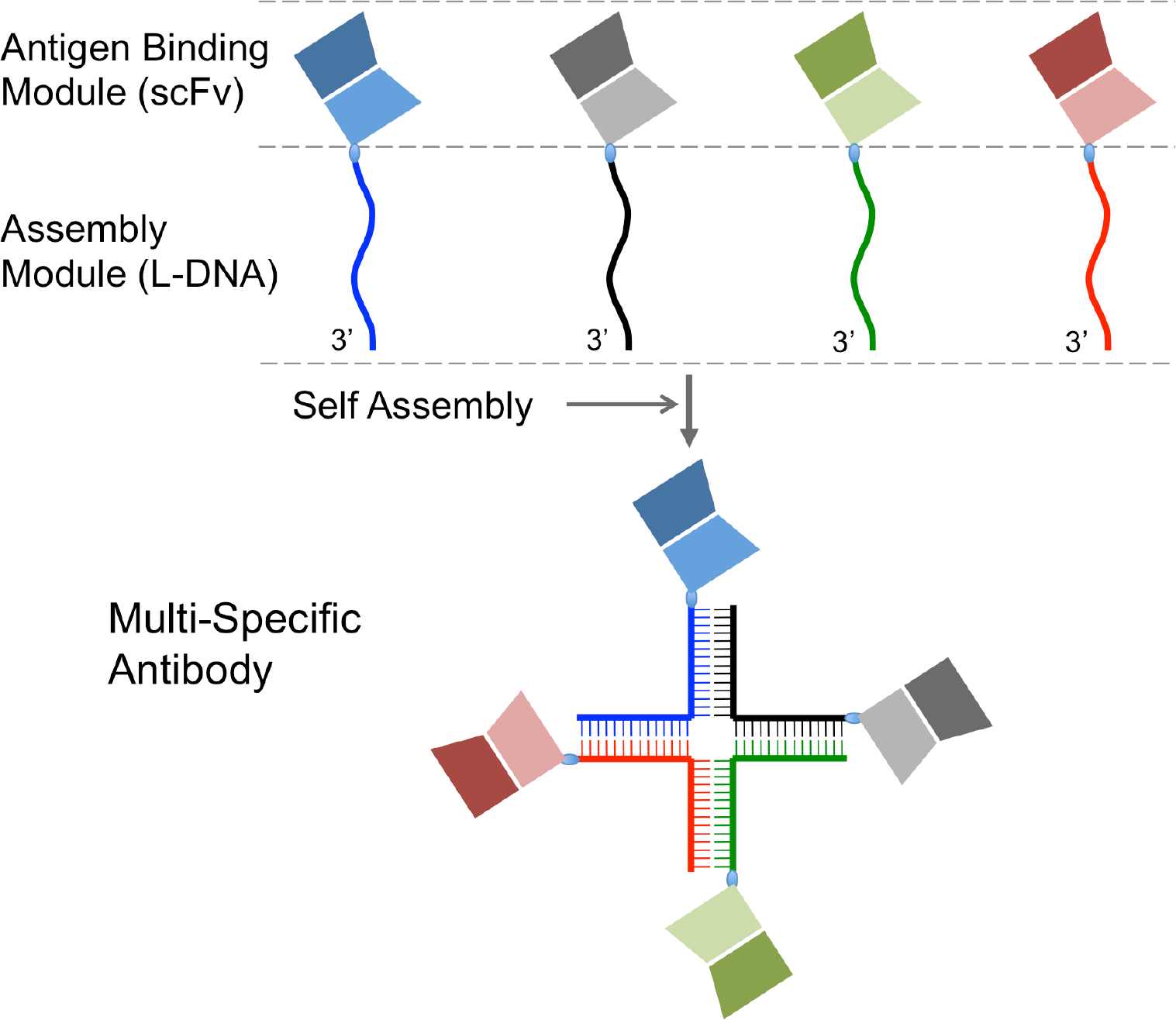
Schematic illustration of the NAPPA implementation for generating multi-specific antibodies.

Several specific details of the above implementation are important for the general applicability of the final assembled product. First, the DNAs should be synthesized as left-handed DNAs (L-DNAs) to prevent degradation by nucleases *in vivo*. L-DNAs are not substrates of any of the known enzymes in nature while still being able to form base-pairing interactions as D-DNAs^17, 18^. Moreover, the L-form nucleic acids are non-immunogenic, as was demonstrated for several L-form aptamers (spiegelmers) that have advanced to clinical trials^19^. Second, the DNA is conjugated to the C-terminus of the scFv so that it is less likely to interfere with the complementarity-determining regions (CDRs). Finally, the length of DNAs should be set such that the melting temperature (*T*_m_) of each of the paired double-stranded segments is greater than 50°C, to ensure the stability of the oligomeric complex. Based on the above considerations, we constructed a MsAb containing two anti-CD19 scFvs (scFv^CD19^) and one anti-CD3 scFv (scFv^CD3^), gathered in a DNA four-way junction complex (**Figure 2a**), designated NAPPA001.

**Figure 2.**
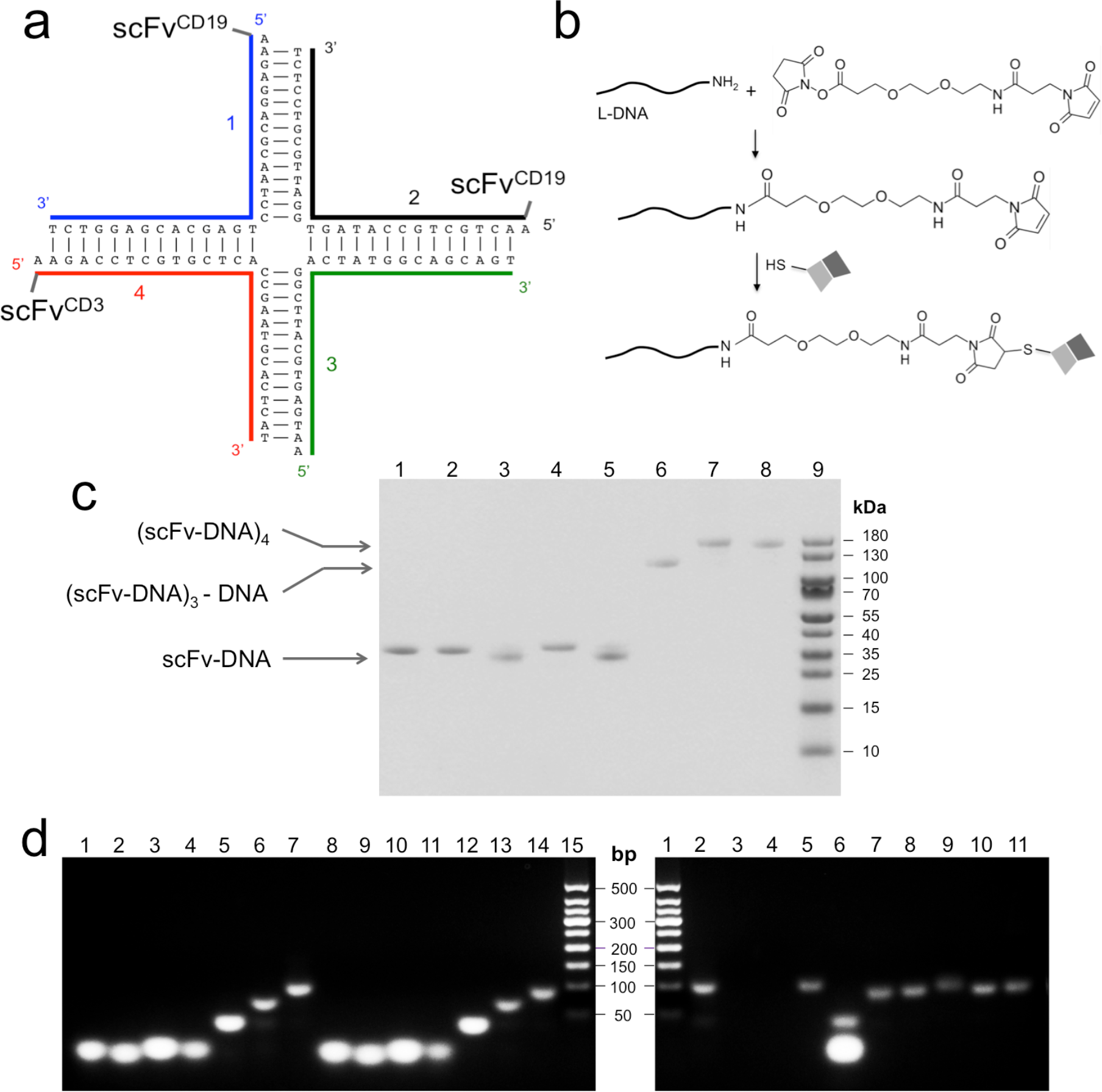
Preparation of the (scFv^CD19^)_2_-scFv^CD3^ antibody using the NAPPA method. (**a**) Sequences of four DNAs that form the 4-way junction and the conjugation of scFv to three of the four DNAs. (**b**) Reaction scheme for linking L-DNA to the scFv-C via PEGylated SMCC. (**c**) SDS-PAGE analysis of various assemblies of scFv-L-DNA conjugates. Lane definitions are: **1** – scFv^CD19^-L-DNA1; **2** – scFv^CD19^-L-DNA2; **3** – scFv^CD3^-L-DNA3; **4** – scFv^4-1BB^-L-DNA4; **5** – scFv^CD3^-L-DNA4; **6** – scFv^CD19^-L-DNA1 + scFv^CD19^-L-DNA2 + L-DNA3 + scFv^CD3^-L-DNA4; **7** – scFv^CD19^-L-DNA1 + scFv^CD19^-L-DNA2 + scFv^CD3^-L-DNA3 + scFv^CD3^-L-DNA4; **8** – scFv^CD19^-L-DNA1 + scFv^CD19^-L-DNA2 + scFv^4-1BB^-L-DNA3 + scFv^CD3^-L-DNA4; **9** – M.W. Marker. (**d**) Agarose gel electrophoresis analysis of L-DNA assemblies and resistance to nucleases. Lanes in the *left* gel: **1** – D-DNA1; **2** – D-DNA2; **3** – D-DNA3; **4** – D-DNA4; **5** – D-DNAs 1, 2; **6** – D-DNAs 1-3; **7** – D-DNAs 1-4; **8-14** – same as 1-7 but for L-DNAs; **15** – MW Marker. Lanes in the *right* gel: **1** – MW Marker; **2** – assembled D-DNA tetramer (1-4); **3-6** – D-DNA tetramer with added DNAase, S1, Exol, and T7 nucleases, respectively; **7** – L-DNA tetramer (1-4); **8-11** – same as in 3-6 but for L-DNA tetramer.

The sequence of the scFv^CD19^ was taken from the CD19-CD3 bispecific antibody Blincyto^®^, which is the first FDA-approved BsAb for use in cancer immunotherapy^10^. The scFv^CD3^ sequence used here is from the anti-CD3 mAb OKT3^20^. The scFvs, all containing a free cysteine at the C-terminus (scFv-C) for DNA conjugation, were expressed in *Escherichia coli* as a C-terminal fusion component to the polyhistidine-tagged maltose binding protein (H_6_-MBP). A TEV cleavage site was inserted between MBP and scFv for separation of the two during purification. The purification of the scFv-DNA conjugates was straightforward (details described in Supporting Information). Briefly, the H_6_-MBP-scFv-C was purified using amylose affinity and treated with TEV enzyme to release strep tagged-scFv-C (scFv-C). The scFv-C was purified by size exclusion chromatography, followed by conjugation to L-DNA at its C-terminal Cys (**Figure 2b**). For chemical conjugation, the 5’ end of the L-DNA was first linked to the PEGylated SMCC bifunctional linker via a reaction of a primary amine on the L-DNA with the NHS ester of the linker. The purified SMCC-PEG2-L-DNA was then linked to the scFv-C via a reaction of the SMCC maleimide with the free thiol of the fusion protein. The reaction mixture was then subject to Strep-Tactin affinity chromatography for removing free DNA. In the final step, ion-exchange chromatography was used to remove proteins not conjugated to DNA, yielding pure scFv-L-DNA. Reducing agent (10 μM TCEP) was used throughout all purification steps before conjugation reaction to keep C-terminal free Cys reduced.

Following the above protocol, we prepared three scFv-L-DNAs: two scFv^CD19^ linked to DNAs 1 and 2, respectively, and one scFv^CD3^ linked to DNA4 (DNA sequences shown in **Figure 2a**). The final MsAb, designated NAPPA001, was assembled by mixing equal moles of scFv^CD19^-L-DNA1, scFv^CD19^-L-DNA2, L-DNA3, and scFv^CD3^-L-DNA4 (**Figure 2c**). Nuclease diggestion tests showed that the assembled L-DNA tetramer was completely resistant to DNAse I, T7 endonuclease, S1 nuclease, and Exonuclease I, whereas that assembled with corresponding D-DNAs was degraded rapidly (**Figure 2d**). We have also shown that the assembled L-DNA tetramer is stable in fetal bovine serum (**Figure S1**).

We next tested the ability of the scFv-L-DNA oligomer to engage T-cells against the Raji cells, which are B-lymphocytes that express CD19 on the cell surface. The CD3-expressing T-cells were from human peripheral blood mononuclear cell (hPBMC). The (scFv^CD19^)_2_-scFv^CD3^ has two ABMs to interact with the Raji cell CD19s and one ABM to interact with the T-cell CD3. Engaging the two cells should activate the T-cells, leading to the formation of immunological synapses with the Raji cells and their eventual lysis. In this experiment, we used [Effector T-cell]:[Tumor cell] (E:T) ratio of 10:1, and examined tumor lysis and T-cell activation induced by various doses of the new MsAb. For tumor lysis, we used LDH (lactate dehydrogenase) detection kit to quantify LDH release from dead cells with impaired membrane integrity. To quantitate T-cell activation, we measured the expression level of CD25 or CD137 (4-1BB) on CD3^+^ cells by flow cytometry analysis of CD25/CD3 or CD137/CD3 co-immunostaining, respectively. The *in vitro* assay showed favorable efficacy of NAPPA001 in inducing tumor cell lysis (**Figure 3a**). Although NAPPA001 was generated from two scFvs with relatively low affinity (*K*_D_ ~10^−7^ M) (**Figure S2**), it showed powerful target cell killing with a EC_50_ of ~0.2 nM after two days of incubation. Independent control experiments have been performed to address the possibility that multivalent antibody binding alone can be toxic to lymphoma cells^21^. These experiments showed that 1) the NAPPA001 does not induce Raji cell lysis in the absence of hPBMC (**Figure S3**), 2) the unassembled scFv^CD19^ and scFv^CD3^ in the absence or presence of L-DNA Holliday junction also do not induce significant cytotoxicity in the presence of hPBMC (**Figure S3**), and 3) scFv^CD19^-scFv^CD3^ BsAb assembled with L-DNA duplex can induce Raji cell lysis in the presence of hPBMC (**Figure S3**), thus confirming that the cytotoxicity was due to T-cell redirecting mediated specifically by the MsAb. In another experiment, the T-cells were activated by the presence of 10 pM NAPPA001 according to the increased expression level of CD25, a biomarker for late stage T-cell activation (**Figure 3b**). Further, cell surface expression of the costimulatory receptor 4-1BB was also upregulated by the MsAb treatment (**Figure 3b**), giving rise to the possibility of further leveraging T-cell activity through simultaneous activation of 4-1BB with our flexible MsAb platform.

**Figure 3.**
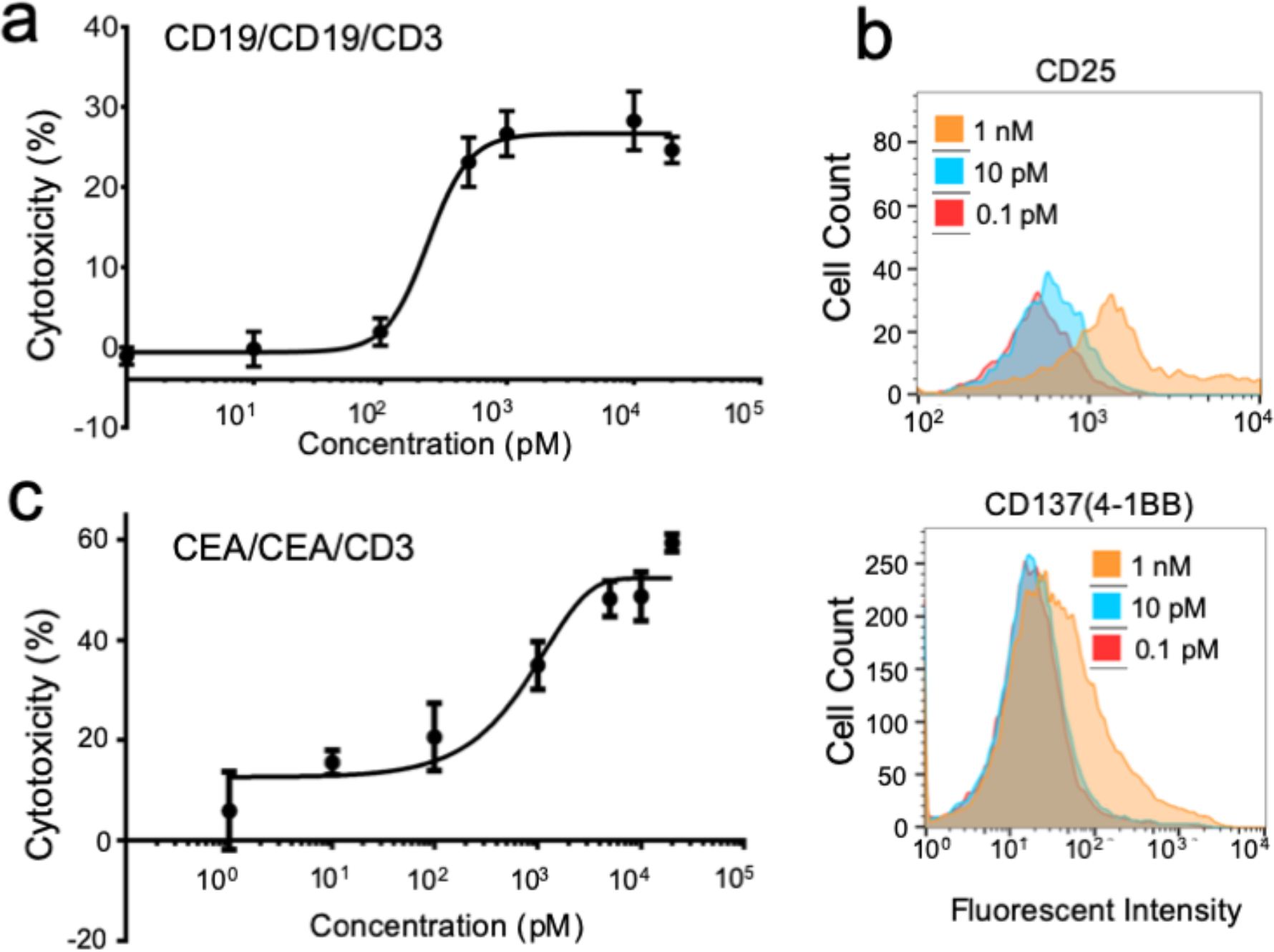
T-cell redirecting function of (scFv^CD19^)_2_-scFv^CD3^ and (scFv^CEA^)_2_-scFv^CD3^. (**a**) Effect of treating CD19^+^ Raji cells with NAPPA001 (at various concentration) and hPBMC (E:T ratio 10:1). Cell viability was measured using the LDH detection kit after 2 days of incubation in 96-well plate. Cytotoxicity (%) was calculated as [sample LDH release – background LDH release]/[maximum LDH release – background LDH release] × 100, where background LDH release is from Raji cells incubated with hPBMC without MsAbs and maximum LDH release is from Raji cells treated with 1% Triton X-100. (**b**) Expression levels of CD25 (left) and CD137 (right) on hPBMC cell surface induced by NAPPA001 in the presence of the Raji cells. hPBMC and Raji cells were mixed at E:T ratio of 10:1, followed by the addition of 0.1 pM, 10 pM, or 1 nM of NAPPA001. After 2-day incubation, cells were co-immunostained with anti-CD25 and anti-CD3 antibodies (for detecting CD25 expression on T-cell surface) or anti-CD137 and anti-CD3 antibodies (for detecting CD137 expression), and examined by flow cytometry. (**c**) Effect of treating CEA^+^ LS174T cells with NAPPA002 (see text) and hPBMC (E:T ratio 10:1). Cell viability was measured as described above.

A similar scFv-L-DNA oligomer was constructed to engage T-cells against the CEA^+^ LS174T cells, which is a model cell line for solid tumor – human colorectal adenocarcinoma. In this case, the NAPPA001 (**Figure 2a**) was modified to specifically target CEA (carcinoembryonic antigen) by replacing the two anti-CD19 scFvs with two anti-CEA scFvs, derived from PR1A3 (a mouse anti-human CEA antibody)^22^. Using the same E:T ratio of 10:1, the new MsAb, (scFv^CEA^)_2_-scFv^CD3^ (designated NAPPA002) showed target cell killing with EC_50_ of ~1.0 nM after two days of incubation (**Figure 3c**).

To test whether the scFv-L-DNA complex can be absorbed into animal bloodstream and distributed to the tumor sites via the circulatory system, we examined its antitumor activity in mice in a subcutaneous CD19^+^ Raji xenograft model. In this study, the cultured Raji cells were cografted with hPBMC at E:T ratio of 4:1. Mixed cells were injected subcutaneously in immunodeficient NSG mice. The (scFv^CD19^)_2_-scFv^CD3^ (given at 10 μg/mouse) or vehicle (PBS) was administered intravenously twice per week for four weeks starting 3 days after tumor/hPBMC co-grafting. The results indicate that mice treated with NAPPA001 almost completely suppressed tumor growth until the end of the study (37 days), in contrast to the rapid tumor growth in the control group (treated with PBS) (**Figure 4a**). Serological examination of the NAPPA001-treated group showed no abnormal levels of aspartate aminotransferase (AST) and alanine aminotransferase (ALT) in comparison to the vehicle group, suggesting the MsAb posed no detectable hepatoxicity (**Figure 4b**). Steady increase of mice body weight further excluded any severe MsAbs-related global toxicity (**Figure S4**).

**Figure 4.**
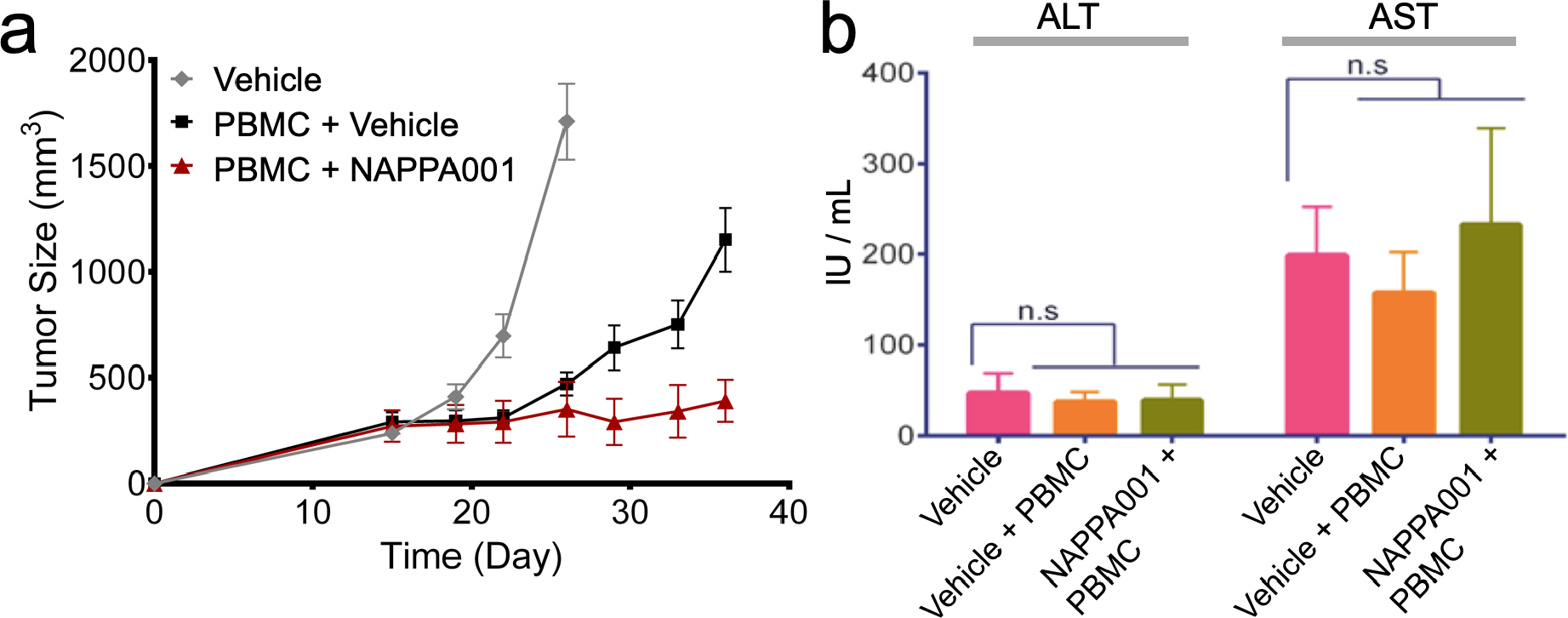
*In vivo* efficacy and toxicity of NAPPA001 in human Burkitt’s lymphoma xenograft mouse model. (**a**) *In vivo* efficacy of NAPPA001 in human Burkitt’s lymphoma xenograft mouse model. Raji tumor cells were implanted or co-implanted with human PBMC (E:T ratio is 4:1) subcutaneously into NSG mice (n = 5) and treated with either PBS (vehicle) in the absence (gray) or presence (black) of PBMC, or PBMC with NAPPA001 (10 μg/mouse) (red) via intravenous injection twice a week starting 3 days after Raji/PBMC co-grafting for 4 weeks. (**b**) Serologic examination of potential hepatotoxicity of NAPPA001. Serum samples were collected from each group 24 hours after the fourth injection of NAPPA001. AST and ALT levels were examined by ELISA in two duplicates, and mean value was calculated from AST and ALT levels in five mice in each group (n = 5).

NAPPA002 ((scFv^CEA^)_2_-scFv^CD3^) was also tested for efficacy in mice using a similar protocol. In this case, CEA^+^ LS174T cells were cografted with hPBMC at E:T ratio of 5:1, and NAPPA002 (given at 20 μg/mouse) or vehicle (PBS) was administered intravenously twice per week starting 3 days from co-grafting for 1.5 weeks. The results show that mice treated with NAPPA002 also significantly suppressed the solid tumor growth compared with the vehicle control groups (**Figure S5**).

We further examined the property of the LS174T-targeting MsAb in targeting and infiltrating solid tumors *in vivo*. For this experiment, we constructed a fluorescent version of NAPPA002 by conjugating the fluorescent probe NHS-Cy5.5 to the 5’ end of L-DNA3 (see **Figure 2a**). The resulting MsAb, (scFv^CEA^)_2_-scFv^CD3^-Cy5.5, was tested in the LS174T xenograft mouse model. After the implanted tumor (in the right flank of mice) reached ~500 mm^3^, (scFv^CEA^)_2_-scFv^CD3^-Cy5.5 or PBS (vehicle) was injected intravenously and the mice were imaged at different time points. We found that (scFv^CEA^)_2_-scFv^CD3^-Cy5.5 rapidly reached the tumor and penetrated into it within one hour, and continued to accumulate at the tumor site, while randomly distributed MsAbs were gradually metabolized (**Figure 5a**). Tumor site was accompanied by strong fluorescence from different directions (ventral and lateral), indicating co-localization of tumor and (scFv^CEA^)_2_-scFv^CD3^-Cy5.5. This result confirmed the ability of the new MsAbs to infiltrate solid tumors. The half-life of (scFv^CEA^)_2_-scFv^CD3^-Cy5.5 was calculated by fluorescence quantification to be ~16 hours (**Figure 5b**).

**Figure 5.**
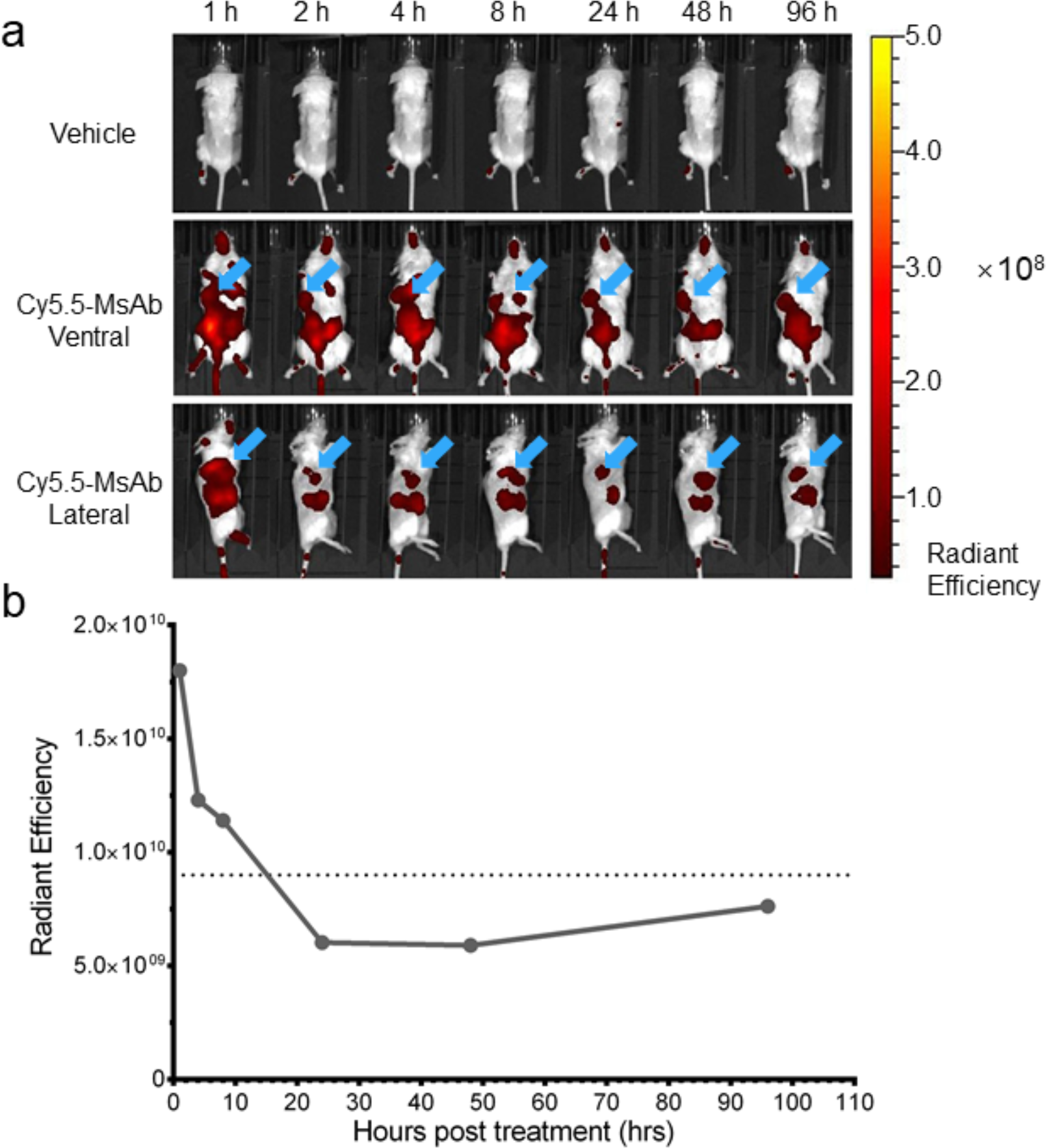
*In vivo* distribution and pharmacokinetics of (scFv^CEA^)_2_-scFv^CD3^-Cy5.5 in LS174T xenograft mouse model. (**a**) Fluorescent images of mice with LS174T and PBMCs (mixed at E:T ratio of 5:1) implanted subcutaneously into the right flanks, injected intravenously with PBS (vehicle) or 100 μg of (scFv^CEA^)_2_-scFv^CD3^-Cy5.5 after the solid tumor reached 500 mm^3^. The fluorescence in mice were monitored and quantified by a small animal imager (PerkinElmer IVIS Kinetic III) at different time points. Fluorescence is represented graphically as radiant efficiency (photons/s/cm^2^/str)/(μW/cm^2^). Blue arrows indicate the position of the implanted tumor. (**b**) Plot of fluorescence intensity at different time points after injection of the fluorescent MsAb, showing that the half-life of the MsAb is ~16 hours.

We have shown that multiple scFvs targeting different antigens, linked together by a L-DNA scaffold, is a stable molecular complex that can efficiently induce T-cell – tumor cell engagement *in vitro* and *in vivo*. We believe the NAPPA format should have little to no toxicity or immunogenicity *in vivo*. Comparing with other MsAb formats, the major addition of NAPPA is the linker-L-DNA. The L-form nucleic acids are non-immunogenic and generally not toxic, since they have advanced to clinical trials^19^. The PEG spacer within the PEGylated SMCC linker should have the effect of enhancing stability and reducing aggregation and immunogenicity^23, 24^. The SMCC linker is widely used in preparing antibody drug conjugate (ADC). For example, in the case of the Trastuzumab-SMCC-DM1 ADC that targets HER2, no SMCC-associated immunogenicity or toxicity was reported in clinical trials^25^.

Probably the biggest advantage of the NAPPA method is that it allows the option to prepare and store individual scFv-L-DNA conjugates separately, each targeting one of the known tumor antigens or immune cell activators, and then assemble a prescribed combination of them into MsAb immediately before use. This type of flexibility would be compatible with personalized cancer immunotherapy, as tumor antigen expression can vary substantially among patients. We found that when the concentrations of the scFv-L-DNA conjugates were determined accurately, mixing them at equal molar ratio could generate very homogeneous oligomeric species.

Another notable advantage is that multi-strand DNA assembly affords unlimited possibilities and one of them is achieving higher order oligomers. We used four complementary DNA strands to generate a BsAb containing three scFvs (two anti-CD19 and one anti-CD3). In this case, two DNA strands carried the same scFv^CD19^, the third did not carry anything, and the fourth carried scFv^CD3^. We note the same set of DNA strands (**Figure 2a**) can be used to generate a tetraspecific antibody that targets the tumor cells more specifically by having, for example, two scFvs that bind to two different tumor surface antigens. Even in the current study, having two anti-CD19 scFvs should theoretically achieve greater specificity for CD19^+^ cells due to avidity effect. Higher order specificity is gaining interest even in therapeutic area outside of immunotherapy. For example, studies have shown that trispecific antibodies targeting three different, highly conserved epitopes of the HIV-1 envelope protein can achieve greater neutralization breadth and potency than single broadly neutralizing antibodies^26, 27^.

Finally, the NAPPA method is, of course, not limited to assembling scFvs. While the assembly modules are nucleic acids, the antigen binding modules can be scFv, nanobody, Fab, or combination of the three. Nanobody, Fab and scFv are generally less stable than IgG, but afford the option of expression in bacteria^28–30^, which is cultured in inexpensive media.

In conclusion, the NAPPA method represents an entirely different means of generating MsAbs. Although the pharmacokinetics and potential toxicity of antibody-nucleic acid conjugates remain to be rigorously tested in higher animals, our current data suggest that the MsAbs assembled by NAPPA can engage T-cells and tumor cells similar to the well-known BsAbs. Therefore, with further optimization, the NAPPA method could potentially be a general solution for generating antibodies with higher order specificity, e.g., trispecific, tetraspecific or higher. Given the ever growing database of effective cancer-specific antigens for immunotherapy, the versatile NAPPA approach potentially also allows rather convenient construction of a library of MsAbs for personalized cancer immunotherapy.

## Experimental Section

Available in Supporting Information

## Supporting Information

Supporting Information is available from the Wiley Online Library or from the author.

## Acknowledgements

This study was supported by internal research fund from the Assembly Medicine, LLC,

## The table of contents entry

This paper reports a *general solution* for generating multi-specific antibodies with *N* degree of specificity. The method is used to generate a multi-specific antibody containing two anti-CD19 and one anti-CD3 antibody fragments (scFvs). The new multi-specific antibody shows strong efficacy in mediating *Raji* tumor cell cytotoxicity in the presence of human T-cells both *in vitro* and *in vivo*.

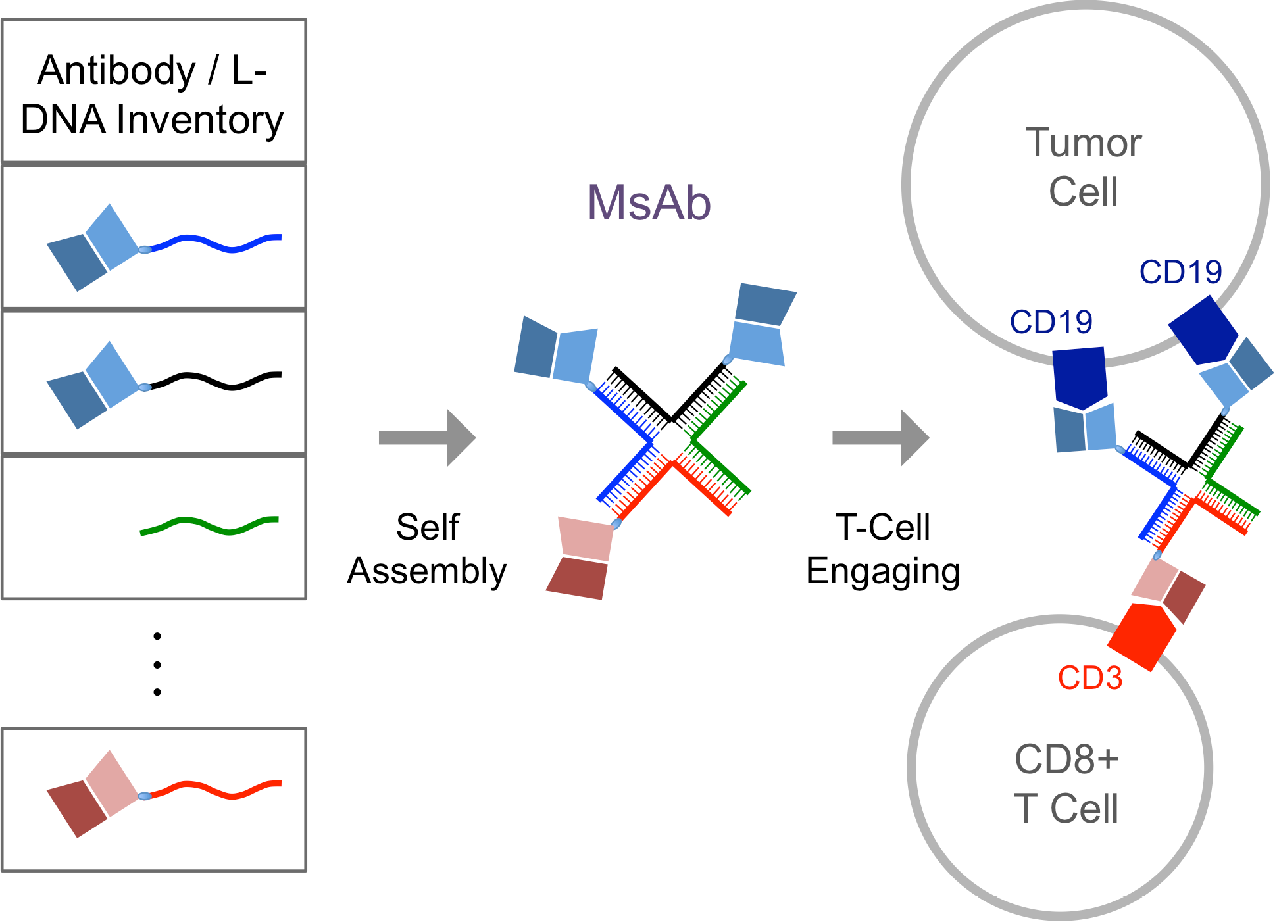

## Literature Cited

[1] U. Brinkmann, R. E. Kontermann. MAbs 2017, 9, 182–212.

[2] Y. Cao, M. R. Suresh. Bioconjug Chem 1998, 9, 635–44.

[3] F. H. Valone, P. A. Kaufman, P. M. Guyre, L. D. Lewis, V. Memoli, Y. Deo, R. Graziano, J. L. Fisher, L. Meyer, M. Mrozek-Orlowski, et al. J Clin Oncol 1995, 13, 2281–92.

[4] A. Thakur, L. G. Lum. Curr Opin Mol Ther 2010, 12, 340–9.

[5] N. D. James, P. J. Atherton, J. Jones, A. J. Howie, S. Tchekmedyian, R. T. Curnow. Br J Cancer 2001, 85, 152–6.

[6] M. G. Fury, A. Lipton, K. M. Smith, C. B. Winston, D. G. Pfister. Cancer Immunol Immunother 2008, 57, 155–63.

[7] T. Nitta, K. Sato, H. Yagita, K. Okumura, S. Ishii. Lancet 1990, 335, 368–71.

[8] W. D. Mallender, E. W. Voss, Jr. J Biol Chem 1994, 269, 199–206.

[9] D. Saerens, G. H. Ghassabeh, S. Muyldermans. Curr Opin Pharmacol 2008, 8, 600–8.

[10] D. Nagorsen, P. Kufer, P. A. Baeuerle, R. Bargou. Pharmacol Ther 2012, 136, 334–42.

[11] E. Wolf, R. Hofmeister, P. Kufer, B. Schlereth, P. A. Baeuerle. Drug Discov Today 2005, 10, 1237–44.

[12] J. B. Ridgway, L. G. Presta, P. Carter. Protein Eng 1996, 9, 617–21.

[13] T. S. Von Kreudenstein, E. Escobar-Carbrera, P. I. Lario, I. D’Angelo, K. Brault, J. Kelly, Y. Durocher, J. Baardsnes, R. J. Woods, M. H. Xie, P. A. Girod, M. D. Suits, M. J. Boulanger, D. K. Poon, G. Y. Ng, S. B. Dixit. MAbs 2013, 5, 646–54.

[14] K. Gunasekaran, M. Pentony, M. Shen, L. Garrett, C. Forte, A. Woodward, S. B. Ng, T. Born, M. Retter, K. Manchulenko, H. Sweet, I. N. Foltz, M. Wittekind, W. Yan. J Biol Chem 2010, 285, 19637–46.

[15] M. Bacac, T. Fauti, J. Sam, S. Colombetti, T. Weinzierl, D. Ouaret, W. Bodmer, S. Lehmann, T. Hofer, R. J. Hosse, E. Moessner, O. Ast, P. Bruenker, S. Grau-Richards, T. Schaller, A. Seidl, C. Gerdes, M. Perro, V. Nicolini, N. Steinhoff, S. Dudal, S. Neumann, T. von Hirschheydt, C. Jaeger, J. Saro, V. Karanikas, C. Klein, P. Umana. Clin Cancer Res 2016, 22, 3286–97.

[16] J. J. Chou, L. Pan, inventorsPreparation of Multi-Specific Protein Drug Library and Its Method of Use. China patent PCT/CN2018/080058. 2017.

[17] K. P. Williams, X. H. Liu, T. N. Schumacher, H. Y. Lin, D. A. Ausiello, P. S. Kim, D. P. Bartel. Proc Natl Acad Sci U S A 1997, 94, 11285–90.

[18] M. J. Damha, P. A. Giannaris, P. Marfey. Biochemistry 1994, 33, 7877–85.

[19] M. Boyce, S. Warrington, B. Cortezi, S. Zollner, S. Vauleon, D. W. Swinkels, L. Summo, F. Schwoebel, K. Riecke. Br J Pharmacol 2016, 173, 1580–8.

[20] J. P. Van Wauwe, J. R. De Mey, J. G. Goossens. J Immunol 1980, 124, 2708–13.

[21] J. Wang, S. Tian, R. A. Petros, M. E. Napier, J. M. Desimone. J Am Chem Soc 2010, 132, 11306–13.

[22] H. Durbin, S. Young, L. M. Stewart, F. Wrba, A. J. Rowan, D. Snary, W. F. Bodmer. Proc Natl Acad Sci U S A 1994, 91, 4313–7.

[23] R. B. Greenwald, Y. H. Choe, J. McGuire, C. D. Conover. Adv Drug Deliv Rev 2003, 55, 217–50.

[24] N. Jain, S. W. Smith, S. Ghone, B. Tomczuk. Pharm Res 2015, 32, 3526–40.

[25] S. Girish, M. Gupta, B. Wang, D. Lu, I. E. Krop, C. L. Vogel, H. A. Burris Iii, P. M. LoRusso, J. H. Yi, O. Saad, B. Tong, Y. W. Chu, S. Holden, A. Joshi. Cancer Chemother Pharmacol 2012, 69, 1229–40.

[26] L. Xu, A. Pegu, E. Rao, N. Doria-Rose, J. Beninga, K. McKee, D. M. Lord, R. R. Wei, G. Deng, M. Louder, S. D. Schmidt, Z. Mankoff, L. Wu, M. Asokan, C. Beil, C. Lange, W. D. Leuschner, J. Kruip, R. Sendak, Y. D. Kwon, T. Zhou, X. Chen, R. T. Bailer, K. Wang, M. Choe, L. J. Tartaglia, D. H. Barouch, S. O’Dell, J. P. Todd, D. R. Burton, M. Roederer, M. Connors, R. A. Koup, P. D. Kwong, Z. Y. Yang, J. R. Mascola, G. J. Nabel. Science 2017, 358, 85–90.

[27] J. J. Steinhardt, J. Guenaga, H. L. Turner, K. McKee, M. K. Louder, S. O’Dell, C. I. Chiang, L. Lei, A. Galkin, A. K. Andrianov, A. D.-R. N, R. T. Bailer, A. B. Ward, J. R. Mascola, Y. Li. Nat Commun 2018, 9, 877.

[28] E. Pardon, T. Laeremans, S. Triest, S. G. Rasmussen, A. Wohlkonig, A. Ruf, S. Muyldermans, W. G. Hol, B. K. Kobilka, J. Steyaert. Nat Protoc 2014, 9, 674–93.

[29] P. Carter, R. F. Kelley, M. L. Rodrigues, B. Snedecor, M. Covarrubias, M. D. Velligan, W. L. Wong, A. M. Rowland, C. E. Kotts, M. E. Carver, et al. Biotechnology (N Y) 1992, 10, 163–7.

[30] K. D. Miller, J. Weaver-Feldhaus, S. A. Gray, R. W. Siegel, M. J. Feldhaus. Protein Expr Purif 2005, 42, 255–67.

