## Supplementary Methods and Figures for "DNA-mediated assembly of multi-specific antibodies for T cell engaging and tumor killing"

### L-DNA preparation

L-DNA amidites (Beta-L-deoxy Adenosine (n-bz) CED phosphoramidite, Beta-L-deoxy Guanosine (n-ibu) CED phosphoramidite, Beta-L-Thymidine CED phosphoramidite and Beta-L-deoxy Cytidine (n-acetyl) CED phosphoramidite) were synthesized by ChemGenes (Massachusetts, USA). Four 5'-modified ($NH_2$-C6) L-DNA fragments (designed to form Holliday junction) were synthesized and purified by Bio-Synthesis (Texas, USA). The four L-DNA fragments are:
L-DNA1 (5'-AAGAGGACGCAATCCTGAGCACGAGGTCT-3'),
L-DNA2 (5'-AACTGCTGCCATAGTGGATTGCGTCCTCT-3'),
L-DNA3 (5'-AATGAGTGCATTCGGACTATCCGAGCAGT-3'), and
L-DNA4 (5'-AAGACCTCGTGCTCACCGAATGCACTCAT-3').

### L-DNA assembly and stability

To test L-DNA assembly, L-DNAs were mixed at equal molar ratio (10 μM each) at room temperature for 1 min, followed by examination with 3% agarose gel. For evaluating the resistance of the L-DNAs to nucleases, four corresponding D-DNAs (with identical sequence to L-DNA) were synthesized and assembled. Assembled L-DNA and D-DNA tetramers were treated at 37 °C for 24 h with different nucleases, including DNase I (Thermo Scientific, AM2222), Exonuclease I (NEB, M0293S), S1 Nuclease (Thermo Scientific, EN0321), and T7 Endonuclease I (NEB, M0302S). The D/L-DNA samples treated with nucleases were examined with 3% agarose gel. To test L-DNA tetramer stability in serum, L-DNA tetramer was mixed with Fetal bovine serum (Gibco) at equal volume ratio (1:1), followed by incubation at 37 °C for 1 h, 4 h, 24 h or 48 h. Samples were examined with 2% agarose gel electrophoresis.

### scFv expression and purification

Gene fragment encoding H6-MBP-(TEV site)-Strep-scFv were subcloned into the pET21(+) vector (Novagen) to generate scFv expression plasmids. Expression constructs were transformed into SHuffle T7 competent *E.coli* cells (New England Biolabs). The cells were grown at 37 °C in LB media until the OD600 reached 0.6~0.8. Then the cell culture was cooled to 16 °C before adding 0.1 mM β-D-thiogalatopyranoside (IPTG), and protein expression lasted overnight at 16 °C. The cells were harvested and resuspended in lysis buffer (20 mM Tris-HCl, 200 mM NaCl，2 mM EDTA, 10 μM TCEP, pH 7.4), followed by sonication with a 6 mm probe (20% output) for 5 min. The cell lysis was centrifuged at 40,000 g for 25 min at 4 °C and the supernatant was loaded to an amylose column (New England Biolabs) pre-equilibrated with the lysis buffer. The fusion protein was eluted with 4 column volumn (CV) of lysis buffer supplemented with 10 mM D-maltose (Sigma). The purified fusion protein was cleaved with tobacco etch virus (TEV) protease (at a concentration of 10 μg/ml) for 2 h at 37 °C to remove the MBP, followed by further purification by size exclusion chromatography (Superdex 200 Increase, GE lifescience). The elution fractions containing the Strep-tagged scFv were collected and stored at -80 °C.

**SPR measurement of scFv affinity to target antigens**

SPR measurements were performed using the Biacore 8K instrument (GE Life Sciences) at 25 °C. Human CD19 ($CD19_{20\text{-}291}$-10xHis, Acro Biosystems, Cat# CD9-H52H2) or CD3 ($CD3_{23\text{-}126}$-6xHis, Abcam, Cat# ab220577) antigen proteins were coupled to the CM5 chip surface by amino coupling. To measure binding, purified MBP-scFvs in the SPR buffer was injected at increasing concentrations (0.0078125, 0.015625, 0.03125, 0.0625, 0.125, 0.25 and 0.5 μM). The response in signal (RU) at each of the above protein concentration was recorded (**Figure S1**). SPR data processing and analysis were performed using the program BIAevaluation (GE Life Sciences). Equilibrium signal ($R_{eq}$) at different MBP-scFv concentration (C) were plotted and fit to the equation $R_{eq} = R_{max}/(1 + K_D/C)$ to determine the equilibrium binding constant ($K_D$). $R_{max}$ is the binding signal at saturation.

**scFv-DNA ligation**

The L-DNA with a 5'-$NH_2$ group was dissolved in PBS (10 mM $Na_2HPO_4$, 1.8 mM $KH_2PO_4$, 2.7 mM KCl, 137 mM NaCl, pH7.4) to a concentration of 0.5 mM. In order to ligate scFv with L-DNA, a mixture of L-DNA and $SM(PEG)_2$ (ThermoFisher) at a molar ratio of 1 : 50 was first incubated at 25 °C for 2 h to ligate $SM(PEG)_2$ with L-DNA. Precipitate formed immediately after adding 2 volumns of anhydrous ethanol. Following incubation at -20 °C for 20 min, the precipitate was collected by centrifuging at 12000 rpm for 10 min and the supernatant was removed. The pellet was washed with 75% ethanol for 4 times. The residual ethanol was vaporized and the pellet was solubilized with double distilled water to a concentration of ~ 5 mg/ml. The $SM(PEG)_2$-DNA was stored at -80 °C.

ScFv was mixed with $SM(PEG)_2$-DNA at a molar ratio of 1:1.2 - 1:2 at 4 °C for overnight to ligate scFv with L-DNA. The conjugation yield was ~90%. Unligated $SM(PEG)_2$-DNA was removed by passing through a strepTactin (Qiagen) column (**Figure S6a**). The elution was further loaded on a HiTrap Q column (GE lifescience) and scFv-DNA was eluted by a 20 CV linear gradient from buffer A (20 mM Tris-HCl, 15 mM NaCl, pH 8.5) to buffer B (20 mM Tris-HCl, 1 M NaCl, pH 8.5). The elution fractions containing scFv-DNA was collected and dialysis against PBS for 1.5 h for twice (**Figure S6b**). The sample was concentrated to a concentration of 10 μM.

**Protein assembly**

The tri-specific antibody with two anti-CD19 units and an anti-CD3 unit were prepared by mixing an equal amount of anti-CD19-HJ1, anti-CD19-HJ2, free HJ3 and anti-CD3-HJ4. The quad-specific antibody with two anti-CD19 units and two anti-CD3 unit were prepared by mixing an equal amount of anti-CD19-HJ1, anti-CD19-HJ2, anti-CD3-HJ3 and anti-CD3-HJ4. The quad-specific antibody with two anti-CD19 units, an anti-41BB units and an anti-CD3 unit were prepared by mixing an equal amount of anti-CD19-HJ1, anti-CD19-HJ2, anti-41BB-HJ3 and anti-CD3-HJ4. All the assemblies were stored at -80 °C.

**Cell cultures**

*Human Burkitt's lymphoma Raji cell line.* Raji cells were cultured in RPMI medium (Gibco, ThermoFisher Scientific) supplemented with 10% Fetal Bovine Serum (Gibco, ThermoFisher Scientific) and 100 U/ml Pen-Strep (GIBCO, ThermoFisher Scientific), in a 37 °C, 5% $CO_2$ incubator (ThermoFisher Scientific).

*Human colorectal adenocarcinoma LS174T cell line.* LS174T cells were maintained in EMEM medium (Gibco, ThermoFisher Scientific) supplemented with 10% Fetal Bovine Serum and 100 U/ml Pen-Strep (GIBCO, ThermoFisher Scientific), in a 37 °C, 5% $CO_2$ incubator (ThermoFisher Scientific).

*Human peripheral blood mononuclear cells (PBMC).* For *in vitro* efficacy study, PBMCs were separated from the whole blood of healthy donors by a density gradient centrifugation method using Ficoll Histopaque. For *in vivo* studies, PBMCs were purchased from appropriate vendors (ASTARTE BIOLOGICS).

**_In vitro_ cytotoxicity of NAPPA001 and NAPPA002**

In each of the 96-well plates, Raji cells ($10^4$) were mixed with fresh human PBMCs ($10^5$) at E:T ratio of 10:1, followed by the addition of various concentrations of $(scFv^{CD19})_2$-$scFv^{CD3}$ (NAPPA001). Raji cell lysis was assessed after 48 hours of MsAb treatment using a LDH detection kit (Beyotime, China). LDH stands for lactate dehydrogenase; it is released into the extracellular media by dead cells due to impaired membrane integrity. Cytotoxicity (%) was calculated as [sample LDH release – background LDH release]/[maximum LDH release – background LDH release] × 100, where background LDH release is from Raji cells incubated with PBMC without MsAbs and maximum LDH release is from Raji cells treated with 1% Triton X-100. All measurements were performed in triplicates. The same protocol was used for other positive and negative control experiments. Exactly the same protocol was also used for measuring *in vitro* cytotoxicity of $(scFv^{CEA})_2$-$scFv^{CD3}$ (NAPPA002) on the LS174T colon carcinoma cells.

**T-cell activation by $(scFv^{CD19})_2$-$scFv^{CD3}$ (NAPPA001)**

Raji cells were seeded in 96-well plate at a density of 10,000 cells/well, and mixed with fresh human PBMCs at E:T ratio of 10:1. $(scFv^{CD19})_2$-$scFv^{CD3}$ After 48 hours treatment with $(scFv^{CD19})_2$-$scFv^{CD3}$ at different concentrations (0.1 pM, 10 pM, 1 nM), cells from each well were divided into two aliquots for measuring cell surface expression of CD25 and CD137, respectively.

For CD25 immunostaining, cells from one aliquot were incubated with Rabbit anti-human CD3ε mAb (1/1000 dilution, Abcam, Cat# ab52959) for 30 min and with Mouse anti-human CD25 mAb (0.5 μg/test, Thermo Fisher scientific, Cat# MA5-12680) for 60 min on ice. After washing, cells were further incubated with Goat anti-Mouse IgG H&L (Alexa Fluor 488) (1/2000 dilution, Abcam, Cat# ab150113) and Donkey anti-Rabbit IgG H&L (Alexa Fluor 647) (1/2000 dilution, Abcam, Cat# ab150075) for 30 min on ice. After washing, the cells were examined by flow cytometry, and $CD3^+$ cells were gated for further evaluation of CD25 expression level.

For CD137 (also known as 4-1BB) immunostaining, cells from the other aliquot were incubated with Rabbit anti-human CD137 mAb (1/400 dilution, Cell Signaling Technology, Cat# 34594) for 1 hour on ice. After washing, cells were further incubated with Mouse anti-human CD3ε mAb [UCHT1] (1/100 dilution, Abcam, Cat# ab34275) and Donkey anti-Rabbit IgG H&L (Alexa Fluor 647) (1/2000 dilution, Abcam, Cat# ab150075) for 30 min on ice. After washing, cells were examined by flow cytometry, and $CD3^+$ cells were gated for further evaluation of CD137 expression level.

**Animal study**

<u>*Mice*.</u> Female (non-pregnant and nulliparous) NOD-*scid* $IL2Rgamma^{null}$ (The Jackson Laboratory, Lot# 005557) mice (age 6-8 weeks) were used for human carcinoma xenograft mouse model. Mice were maintained under pathogen-free and standardized environmental conditions (20±1 °C, 50±10% relative humidity, 12 hours light-dark daily cycles). All

experiments were done according to the American Animal Protection Law with permission from the responsible local authorities.

_In vivo efficacy of $(scFv^{CD19})_2$-$scFv^{CD3}$ (NAPPA001)._ Raji cells ($5\times10^5$ cells) (50 μL) were pre-mixed with human PBMC ($2\times10^6$ cells) (50 μL) at an E:T ratio of 4:1, and further mixed with Matrigel (100 μL) in a 1:1 ratio yielding 200 μL suspension for subcutaneous implantation into the right flanks of NOD/SCID mice. The $(scFv^{CD19})_2$-$scFv^{CD3}$ (0.5 mg per 1 kg of body mass) was injected intravenously twice a week for four weeks starting 3 days after Raji/PBMC co-grafting. Caliper tumor volume measurements were made twice-weekly on all surviving animals. Tumor volumes were estimated by the following formula: Tumor Volume = $0.5 \times (\text{length} \times \text{width}^2)$, where length is the longest diameter measurement and width is the shortest diameter measurement.

_In vivo efficacy of $(scFv^{CEA})_2$-$scFv^{CD3}$ (NAPPA002)._ LS174T cells ($10^6$) were pre-mixed with PBMCs ($5\times10^6$) at an E:T ratio of 5:1, and further mixed with Matrigel in a 1:1 ratio yielding 200 μL suspension for subcutaneous implantation into the right flanks of NOD/SCID mice. The $(scFv^{CEA})_2$-$scFv^{CD3}$ (1 mg per 1 kg of body mass) was injected intravenously twice a week for 1.5 weeks starting 3 days after LS174T/PBMC co-grafting. Caliper tumor volume measurements were made twice-weekly on all surviving animals. Tumor volumes were estimated by the following formula: Tumor Volume = $0.5 \times (\text{length} \times \text{width}^2)$, where length is the longest diameter measurement and width is the shortest diameter measurement.

_In vivo imaging of MsAbs targeting solid tumor._ LS174T cells ($10^6$) were pre-mixed with PBMCs ($5\times10^6$) at an E:T ratio of 5:1, and further mixed with Matrigel in a 1:1 ratio yielding 200 μL suspension for subcutaneous implantation into the right flanks of NOD/SCID mice. When tumor volume reached ~500 $mm^3$, $(scFv^{CEA})_2$-$scFv^{CD3}$-Cy5.5 (220 μL, 0.45 mg/ml) or PBS (vehicle) was injected intravenously. Fluorescence intensity in mice was recorded and quantified by a small animal imager (PerkinElmer IVIS Kinetic III) at different time points (1, 2, 4, 8, 24, 48, 96 h). Fluorescence intensity vs. Time curve was generated to determine half-life of the fluorescent MsAb in mice.

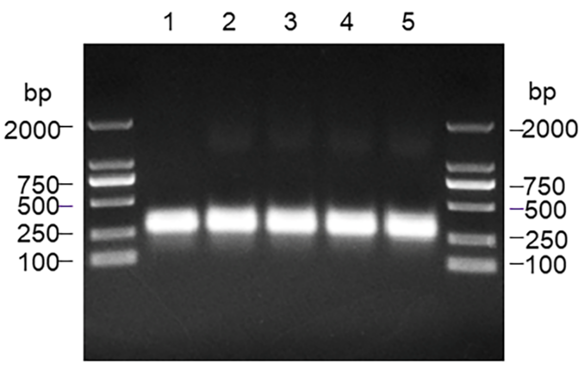


Lane 1 L-tetramer
Lane 2 L-tetramer + Serum, 1h
Lane 3 L-tetramer + Serum, 4h
Lane 4 L-tetramer + Serum, 24h
Lane 5 L-tetramer + Serum, 48h

**Figure S1**. Stability of L-DNA tetramer in serum.

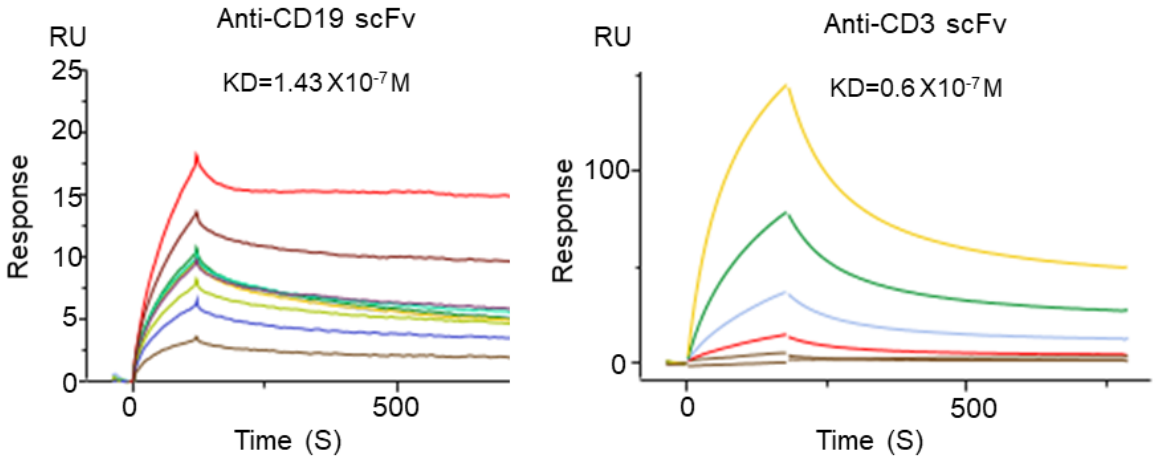


**Figure S2.** Determination of scFv affinity by Surface Plasmon Resonance (SPR).

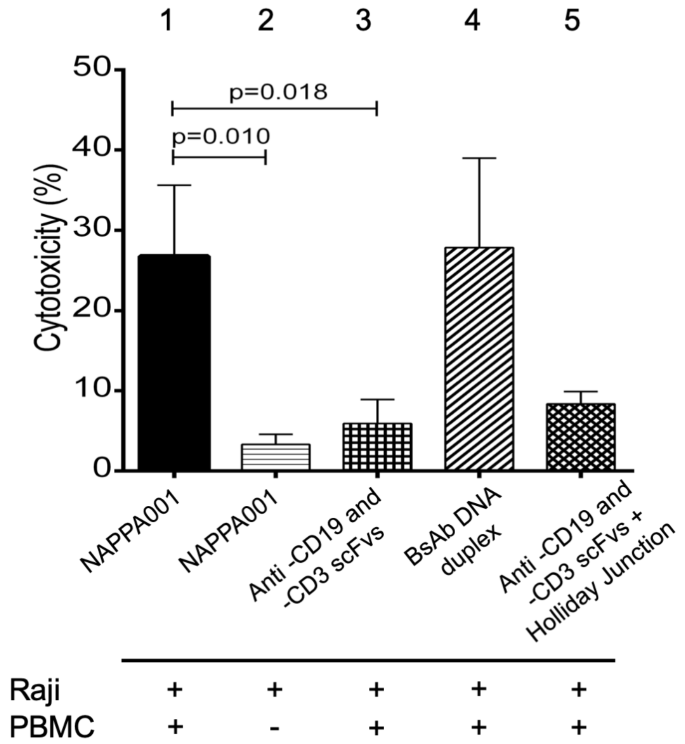


**Figure S3**. Negative and positive controls for T-cell redirected killing of Raji cells mediated by NAPPA001. To examine whether cancer cell killing was due to T-cell redirecting, Raji cells were treated with 500 pM NAPPA001 in the presence (**1**) or absence (**2**) of hPBMC (E:T=10:1) for 2 days. To examine whether the cancer cell killing on the left was due to T-cell − Raji engaging mediated by the bispecific NAPPA001, Raji cells were treated with a mixture of $scFv^{CD19}$ and $scFv^{CD3}$ (500 pM) in the presence of hPBMC (E:T=10:1) (**3**). To test whether $scFv^{CD19}$-$scFv^{CD3}$ assembled with a simple L-DNA duplex can function as a normal BsAb, $scFv^{CD19}$-L-DNA1 and $scFv^{CD3}$-L-DNA4 were mixed at 1:1 ratio. Raji cells were treated with 500 pM of BsAb in the presence of hPBMC (E:T=10:1) for 2 days (**4**). To ensure that antibody assembly was mediated by L-DNA oligomerization, the $scFv^{CD19}$ and $scFv^{CD3}$ were mixed with L-DNA Holliday junction at 500 pM each and used to treat Raji cells in the presence of hPBMC (E:T=10:1) (**5**). Tumor lysis (cytotoxicity) was measured using the LDH (lactate dehydrogenase) detection kit, which quantifies LDH release from dead cells with impaired membrane integrity.

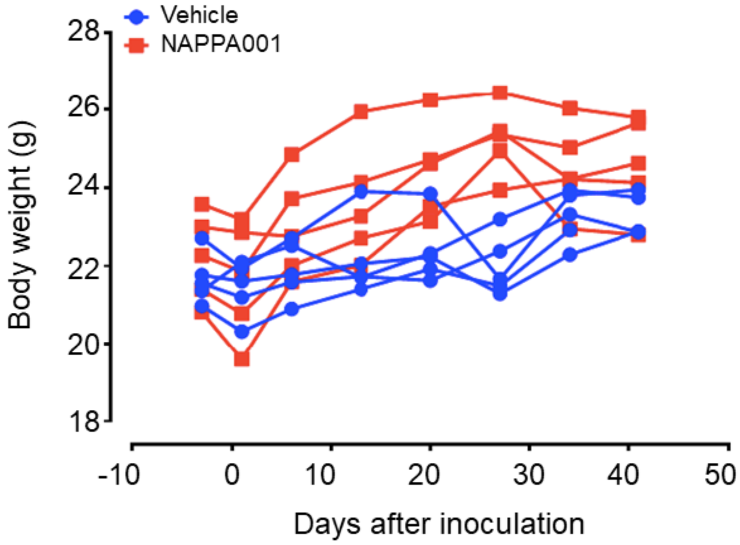


**Figure S4.** Body weights of the five mice during *in vivo* efficacy study of $(scFv^{CD19})_2$-$scFv^{CD3}$ (NAPPA001).

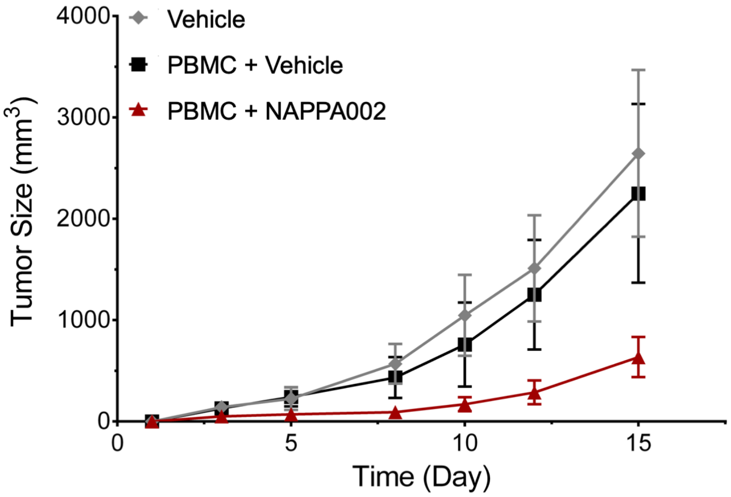


**Figure S5.** *In vivo* efficacy of NAPPA002 in human colorectal adenocarcinoma (LS174T cell line) xenograft mouse model. LS174T tumor cells were implanted or co-implanted with human PBMC (E:T ratio is 5:1) subcutaneously into NSG mice (n = 6) and treated with either PBS (vehicle) in the absence (gray) or presence (black) of PBMC, or NAPPA002 (20 µg/mouse) (red) via intravenous injection twice a week starting 3 days after LS174T/PBMC co-grafting for 1.5 weeks.

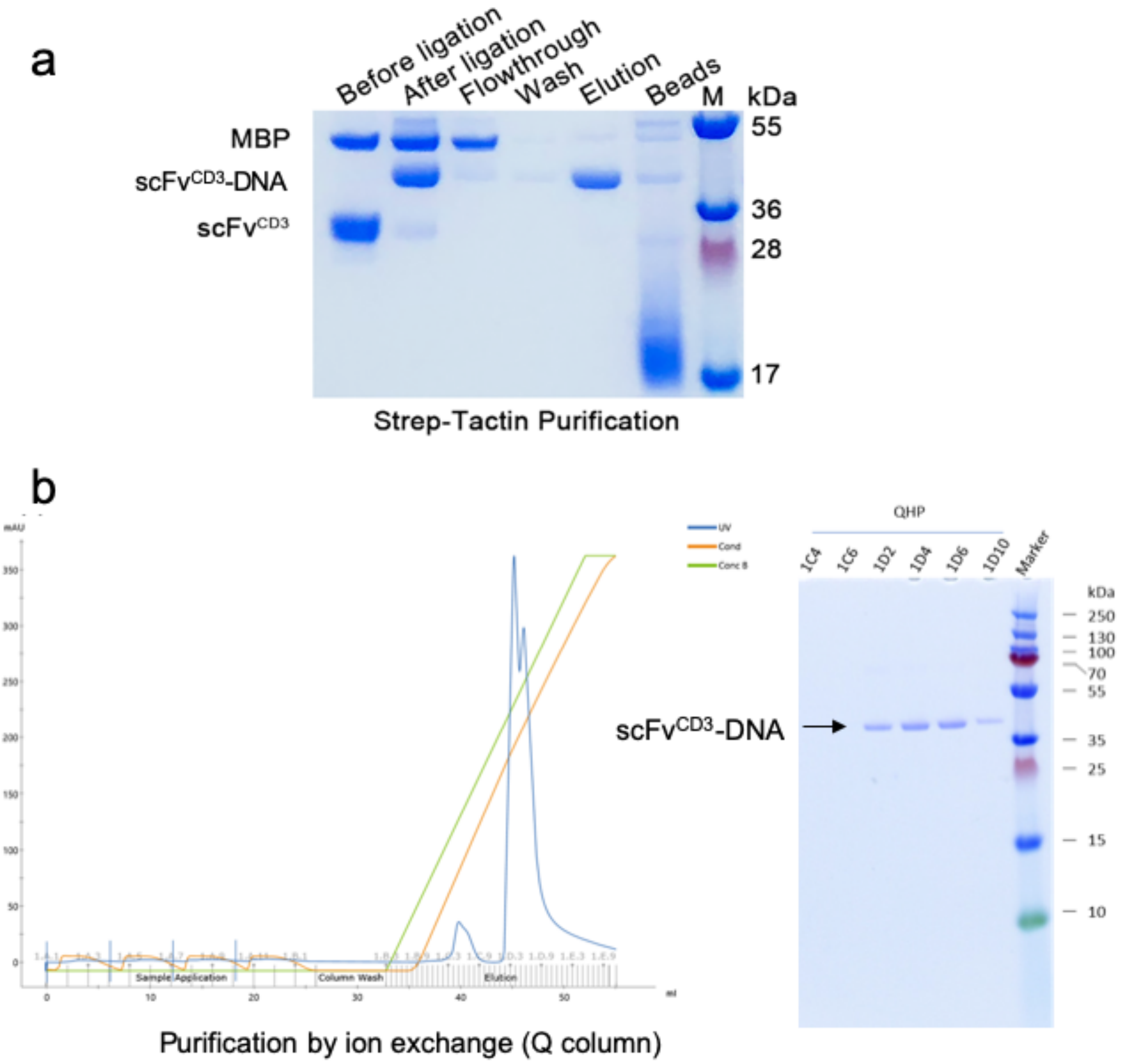


**Figure S6**. Preparation and purification of scFv-DNA. The scFv released from the MBP fusion protein was mixed with SM(PEG)$_2$-L-DNA at a molar ratio of 1:1.2 - 1:2 at 4 °C for overnight to ligate scFv with L-DNA. (**a**) SDS-PAGE analysis of scFv-DNA ligation and removal of unligated DNA by passing through a strepTactin (Qiagen) column. The conjugation yield was ~90%. (**b**) Ion exchange purification of the sample from (a) to remove unligated protein. Elution fractions from the HiTrap Q column (GE lifescience) were analyzed by SDS-PAGE.
